# GroovDB: A database of ligand-inducible transcription factors

**DOI:** 10.1101/2022.07.18.500503

**Authors:** Simon d’Oelsnitz, Andrew D. Ellington

## Abstract

Genetic biosensors are integral to synthetic biology. In particular, ligand-inducible prokaryotic transcription factors are frequently used in high-throughput screening, for dynamic feedback regulation, as multi-layer logic gates, and in diagnostic applications. In order to provide a curated source that users can rely on for engineering applications, we have developed GroovDB (available at https://groov.bio), a Web-accessible database of ligand-inducible transcription factors that contains all information necessary to build chemically-responsive genetic circuits, including biosensor sequence, ligand, and operator data. Ligand and DNA interaction data has been verified against the literature, while an automated data curation pipeline is used to programmatically fetch metadata, structural information, and references for every entry. A custom tool to visualize the natural genetic context of biosensor entries provides additional information that provides potential insights into alternative ligands and systems biology.

## Introduction

The inputs for regulated synthetic circuits are often small molecules, and the dominant class of sensors for such molecules are currently ligand-inducible prokaryotic transcription factors, such as TetR and LacI. While early genetic circuits were generally inducer-agnostic^1,2^, modern circuits are frequently built around the inducer molecules themselves, including circuits for high-throughput screens and diagnostic applications. Synthetic biologists typically ‘mine’ biosensors from the literature^3,4^, by searching through genome databases ^5,6^, or via directed evolution ^7,8^.

Since identifying an appropriate sensor for a given project remains a challenge, we have developed GroovDB, a comprehensive database of inducible transcription factors for synthetic biology and other applications. It currently documents over 100 genetic biosensors from 62 different organisms that can collectively access 131 unique ligands, and can be further expanded by user participation. Database content is organized and managed with a SQL schema supported by a Python-based backend API, and is freely accessible via a modern web application developed with React. Each biosensor entry contains information on ligand interactions, DNA interactions, structures (where available), and associated references. A custom tool displays the natural genetic context of each transcription factor, providing potential insights into alternative inducer molecules. Overall, GroovDB aims to serve as a central resource for synthetic biologists and biotechnologists to facilitate the development of ligand-inducible genetic circuits.

## Results

### Data curation workflow

Biosensor entries are sourced from peer-reviewed research articles (**Figure 1**). A custom text mining application was developed to pre-select publications for further review. The application uses the NCBI’s Entrez API to search for articles to identify PubMed IDs relating to known biosensors (such as “TetR”), and then further searches for keywords such as “EMSA” or “DNase footprinting” (since these are the most common experimental techniques used to identify protein:DNA and/or protein:ligand interactions) using the Beautiful Soup HTML parser. The RegPrecise and PRODORIC databases were also screened for biosensor entries that had associated small molecule ligands, and their corresponding references were collected for review^9,10^.

**Figure 1:**
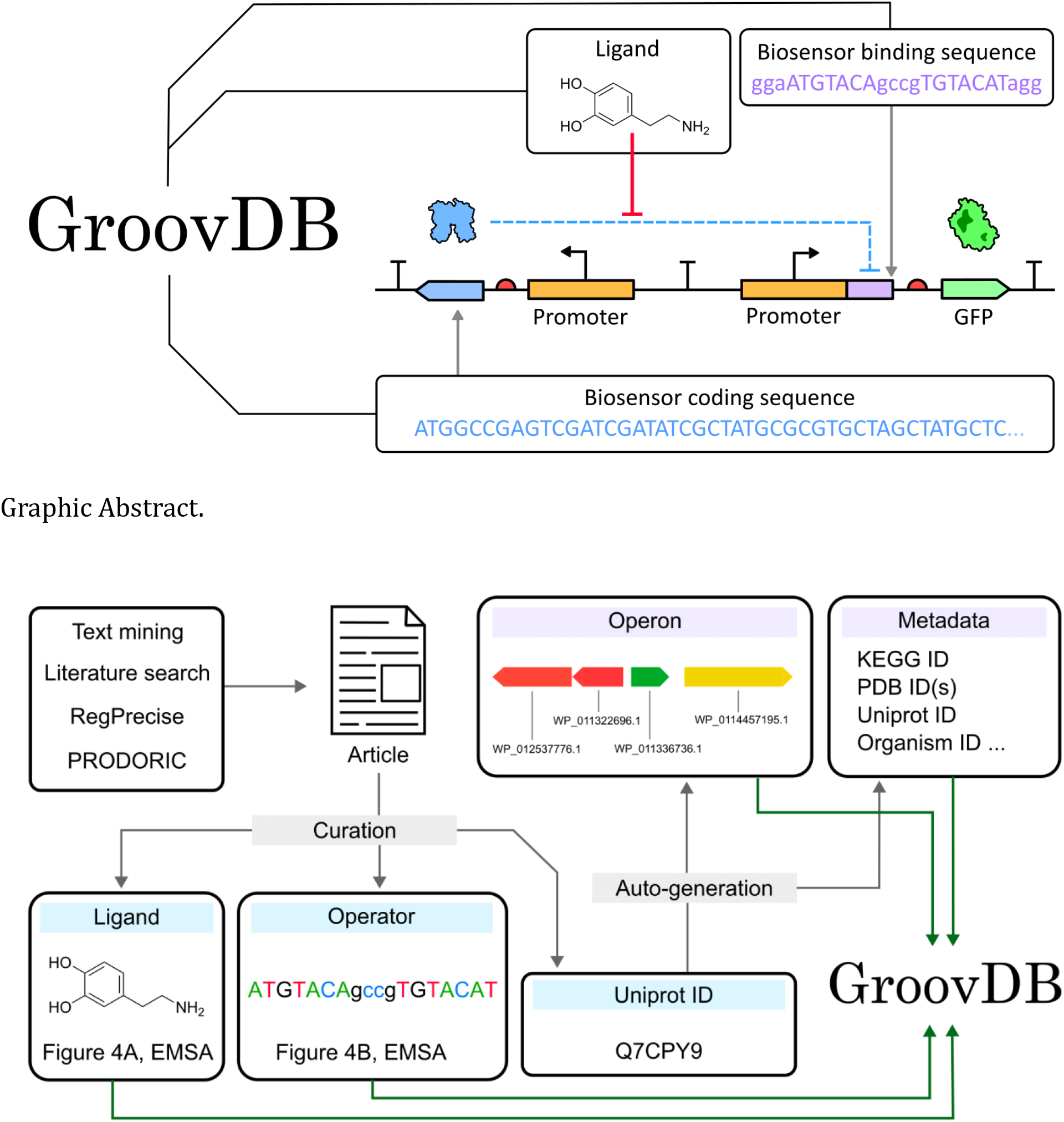
Data curation workflow A custom text-mining program and existing databases are used to curate a list of peer-reviewed research articles from which the Uniprot ID, ligand interaction, and DNA interaction data are extracted for each sensor. The Uniprot ID is then used to programmatically fetch metadata on the biosensor (including the Kegg ID, Uniprot ID, Organism ID, and PDB IDs) in addition to the local genetic context of the biosensor. All of this data is then used to populate a SQL database within GroovDB.

**Figure 2:**
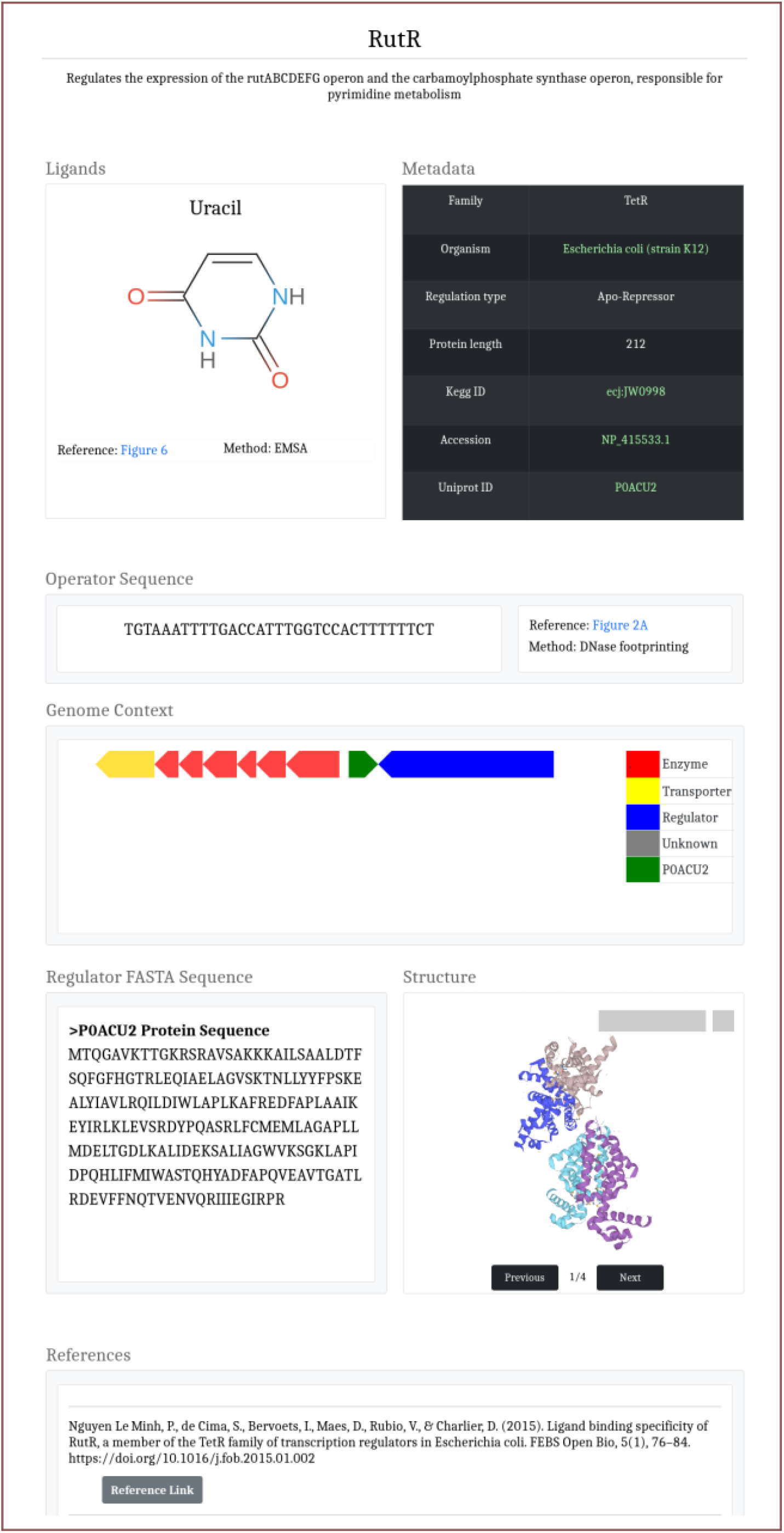
Web interface for GroovDB’s Sensor page.

Protein:ligand interaction data is carefully curated in GroovDB in order to reduce the risk of genetic circuit failure. Acceptable experimental techniques for the inclusion of ligand interactions include electrophoretic mobility shift assays (EMSA), DNase footprinting, surface plasmon resonance, isothermal titration calorimetry, or synthetic regulation in a non-native host (defined as the modulated production of a heterologous protein expressed via the transcription factor’s binding site). We exclude crystal structures of the biosensor in complex with a ligand as sufficient evidence of a valid protein:ligand interaction, since structures of transcription factors have been solved in complex with non-inducer ligands^11,12^. By relying on observations of synthetic regulation in a non-native host we avoid mis-identifying the input due to metabolism of the input, a discrepancy that has been observed in several pathways where cofactor-ligated analogs of an input proved to be the true inducers of a transcription factor being studied^12–14^.

Following identification of suitable biosensors, protein:ligand interactions, protein:DNA interactions, and a Uniprot ID were extracted. The Uniprot ID was subsequently used to extract metadata, including the host organism, RefSeq ID, KEGG ID, any associated PDB IDs, and relevant references. A separate program fetches data on genes that lie within the local genetic context of the biosensor, ultimately outputting a JSON object containing information on the directions, locations, and annotations of neighboring genes. It should be noted that while GroovDB faithfully documents known protein:ligand interactions, sensors may have a range of ligands that they are responsive to; the genetic context of the biosensor (which GroovDB displays, see below) may assist in identifying alternative ligands.

### Database structure

All data is stored in a SQLite database, which provides an organized and easily searchable data structure. Biosensors are linked to ligands and operators via many-to-many relationships, providing flexibility to create complex shared associations between biosensors, ligands, and operators. Data stored in the SQLite database is accessed by a Flask backend API, which processes and passes the data to a React frontend for visualization. Ligands are stored as SMILES strings and protein structures are stored as PDB ID codes to minimize file sizes.

### Data visualization

Biosensors are grouped by their structure-defined family and can be accessed either by text searches on the “Home” page or via browsing through biosensor families on the “Browse” page. For example, GroovDB contains at least 10 entries for the TetR, LysR, AraC, MarR, GntR, LacI, LuxR, and IclR families. Other families with fewer than 10 members, such as TrpR and ROK families, are currently categorized in a separate group.

Once a biosensor entry is selected, the inducer molecule, operator sequence, and biosensor protein sequence are displayed; these are the most important three components necessary to build a regulated genetic circuit. In addition, other modules are presented to provide additional context for circuit engineering, including the biosensor regulatory mechanism, host organism, and protein length, as well as links to external databases (included in a metadata table). If the biosensor’s structure is available, an interactive structure visualization plugin can illustrate key residues involved in ligand recognition. Finally, the genetic context – the set of genes neighboring the biosensor – is displayed, color-coded according to annotated gene functions, and linked to corresponding information in the NCBI protein database. This biology-focused tool is a unique feature of GroovDB, and should help researchers better hypothesize alternative inducer molecules, regulatory targets, or systems biology. For example, if a neighboring regulated gene is an efflux pump known to transport structurally divergent molecules, the corresponding biosensor may also recognize diverse molecules beyond what it has so far been characterized to bind. Similarly, other regulatory genes surrounding the biosensor may indicate ties to the systems biology of the organism and thus can serve to indicate potential metabolic cross-talk and provide clues for further engineering the biosensor in a new context.

### Expanding GroovDB

While we anticipate curating GrooveDB well into the future, its true utility will come with expanded community use. To add new biosensors, a downloadable .xlsx file template (available at https://groov.bio/contact) is provided that includes data fields necessary to populate a sensor entry, as well as an example of valid input data for a biosensor. We have in turn developed an automated workflow that parses data from the .xlsx file and passes it to a data curation workflow (**Figure 1**), thereby automatically creating new SQL biosensor entries for periodic review.

## Discussion

Currently, there exist few resources for identifying genetic biosensors. Several databases, such as MIST, P2CS, and KEGG, provide a comprehensive overview of signal transduction pathways in various organisms, but lack the DNA and ligand binding data of corresponding biosensors necessary for their composition in genetic circuits^15–17^. Other databases, including RegulonDB, DBTBS, CoryneRegNet, and RhizoRegNet, include protein:DNA interactions, but only for biosensors found in specific organisms^18–21^. Finally, the RegPrecise and PRODORIC databases include both protein:DNA and protein:ligand interactions, but RegPrecise is out of date and in PRODORIC fewer than 30% of entries have an associated ligand. Moreover, the criteria for including protein:ligand interactions is not described, and regulator entries cannot be queried based on the cognate ligand^9,10^.

GroovDB provides an updated, community-based, and flexible platform for cataloging the rapidly growing collection of genetic biosensors available to synthetic biologists, molecular biologists, and biotechnologists. We have already successfully used the database to build functional ligand-inducible genetic circuits. For example, by placing GroovDB-sourced operator sequences for the CamR and RamR repressors downstream of the -10 site of a strong *E. coli* promoter driving GFP, we were able to build reporter circuits under control of CamR and RamR biosensors that were responsive to camphor and tetrahydropapaverine, respectively^4,7^. In general, activator-regulated sensor circuits can be generated by placing GroovDB-annotated binding sequences upstream of reporters (such as an RBS and GFP gene), and this approach has been used to successfully build circuits responsive to erythritol with EryD, 3-hydroxypropionate with HpdR, and cis-cis muconate with BenM, among others^3,22,23^. These general design strategies are described in more detail within the documentation section of GroovDB https://groov.bio/howToUse.

## Methods

The Beautiful Soup package was used as a component of the text mining application for parsing HTML files. The SmilesDrawer Javascript component is used to render SMILES strings from the SQL database as 2D chemical structure images^24,25^. The LiteMol plugin is used to display interactive 3D structures of the subject sensor, if available.

## Supporting information

Database Raw Data

## Acknowledgements

This work was supported by grants from the Air Force Office of Scientific Research (FA9550-14-1-0089), National Institutes of Health (5R01EB026533-04), and the Welch Foundation (F-1654). The authors would like to thank Danny Diaz for his contribution to the deployment of GroovDB and Lakshmi Surada for the development of the text mining application.

## Notes

S. D. is the founder of Retna Bio, a company that commercializes genetic biosensors. GroovDB is accessible at https://groov.bio. The text-mining application is available on GitHub at https://github.com/simonsnitz/text-mining

