## Supplementary Table 1 for "GroovDB: A database of ligand-inducible transcription factors"

**Supporting Information**

Supporting Tables S1

**Supporting Table S1: GroovDB Summary Statistics**

| Property | Value |
|---|---|
| Total sensors | 101 |
| Total unique ligands | 131 |
| Total unique organisms | 62 |
| Total unique families | 11 |
| Total Apo-Repressors | 63 |
| Total Co-Repressors | 3 |
| Total Co-Activators | 34 |
| Total Apo-Activators | 1 |
| Entries with structures solved | 34 |
